# Connectivity map of bipolar cells and photoreceptors in the mouse retina

**DOI:** 10.1101/065722

**Authors:** Christian Behrens, Timm Schubert, Silke Haverkamp, Thomas Euler, Philipp Berens

## Abstract

Visual processing begins at the first synapse of the visual system. In the mouse retina, three different types of photoreceptors provide input to 14 bipolar cell (BC) types. Classically, most BC types are thought to contact all cones within their dendritic field; ON BCs would contact cones exclusively via so-called invaginating synapses, while OFF BCs would form basal synapses. By mining publically available electron microscopy data, we discovered interesting violations of these rules of outer retinal connectivity: ON BC type X contacted only ~20% of the cones in its dendritic field and made mostly atypical non-invaginating contacts. Types 5T, 5O and 8 also contacted fewer cones than expected. In addition, we found that rod BCs received input from cones, providing anatomical evidence that rod and cone pathways are interconnected in both directions. This suggests that the organization of the outer plexiform layer is more complex than classically thought.

## Introduction

Visual processing already starts at the very first synapse of the visual system, where photoreceptors distribute the visual signal onto multiple types of bipolar cells. For example, in the mouse retina, two types of cone photoreceptors differing in their spectral properties–short (S-) and medium wavelength-sensitive (M-) cones – and rod photoreceptors provide input to 14 different bipolar cell types (reviewed in Euler et al., 2014). The precise connectivity rules between photoreceptors and bipolar cell (BC) types determine which 39 signals are available to specific downstream circuits. Therefore, the connectome of the outer 40 retina is essential for a complete picture of visual processing in the retina.

For some mouse BC types, specific connectivity patterns have already been described: For 42 example, based on electrical recordings and immunohistochemistry cone bipolar cell type 1 (CBC1) have been suggested to contact selectively M-cones, whereas CBC9 exclusively 44 contacts S-cones (Haverkamp et al., 2005; Breuninger et al., 2011). The other BC types are 45 thought to contact all M-cones within their dendritic field, but the connectivity to S-cones is 46 unclear (Wässle et al., 2009). In addition, two fundamental cone-BC contact shapes have 47 been described: invaginating contacts with the dendritic tips extending into the cone pedicle 48 and flat (basal) contacts that touch the cone pedicle base, commonly associated with ON and OFF-BCs, respectively (Dowling and Boycott, 1966; Kolb, 1970; Hopkins and Boycott, 1995).

Rod bipolar cells (RBCs) are commonly thought to receive exclusively rod input and to feed this signal into the cone pathway via AII amacrine cells (reviewed by Bloomfield and Dacheux, 2001). However, physiological data indicate that RBCs may receive cone photoreceptor input as well (Pang et al., 2010). Also, types CBC3A, CBC3A and CBC4 have also been reported to receive direct rod input (Mataruga et al., 2007; Haverkamp et al., 2008; Tsukamoto and Omi, 2014) suggesting that rod and cone pathways are much more interconnected than their names suggest.

Here we analyzed an existing electron microscopy dataset (Helmstaedter et al., 2013) to quantify the connectivity between photoreceptors and bipolar cells. We did not find evidence for additional M-or S-cone selective CBC types in addition to the reported CBC1 and 9. However, we found interesting violations of established rules of outer retinal connectivity: The newly discovered CBCX (Helmstaedter et al., 2013), likely an ON-CBC (Ichinose et al., 2014), had unexpectedly few and mostly atypical basal contacts to cones. CBC5T, CBC5O and CBC8 also contacted fewer cones than expected from their dendritic field. In addition, 65 we provide anatomical evidence that rod and cone pathways are connected in both 66 directions: Not only OFF-types CBC3A, CBC3B and CBC4 get direct input from rods but also 67 RBCs from cones.

## Results

### Identification of S-and M-cones

We used the serial block-face electron microscopy (SBEM) dataset *e2006* published by Helmstaedter et al. (2013a) to analyze the connectivity between photoreceptors and bipolar cells in the outer plexiform layer (OPL) of the mouse retina (Figure 1A). To this end, we reconstructed the volume of all cone axon terminals (cone pedicles; n=163) in the dataset as well as the dendritic trees of all BCs (n=451; Figure 1B, see *Methods*).

**Figure 1:**
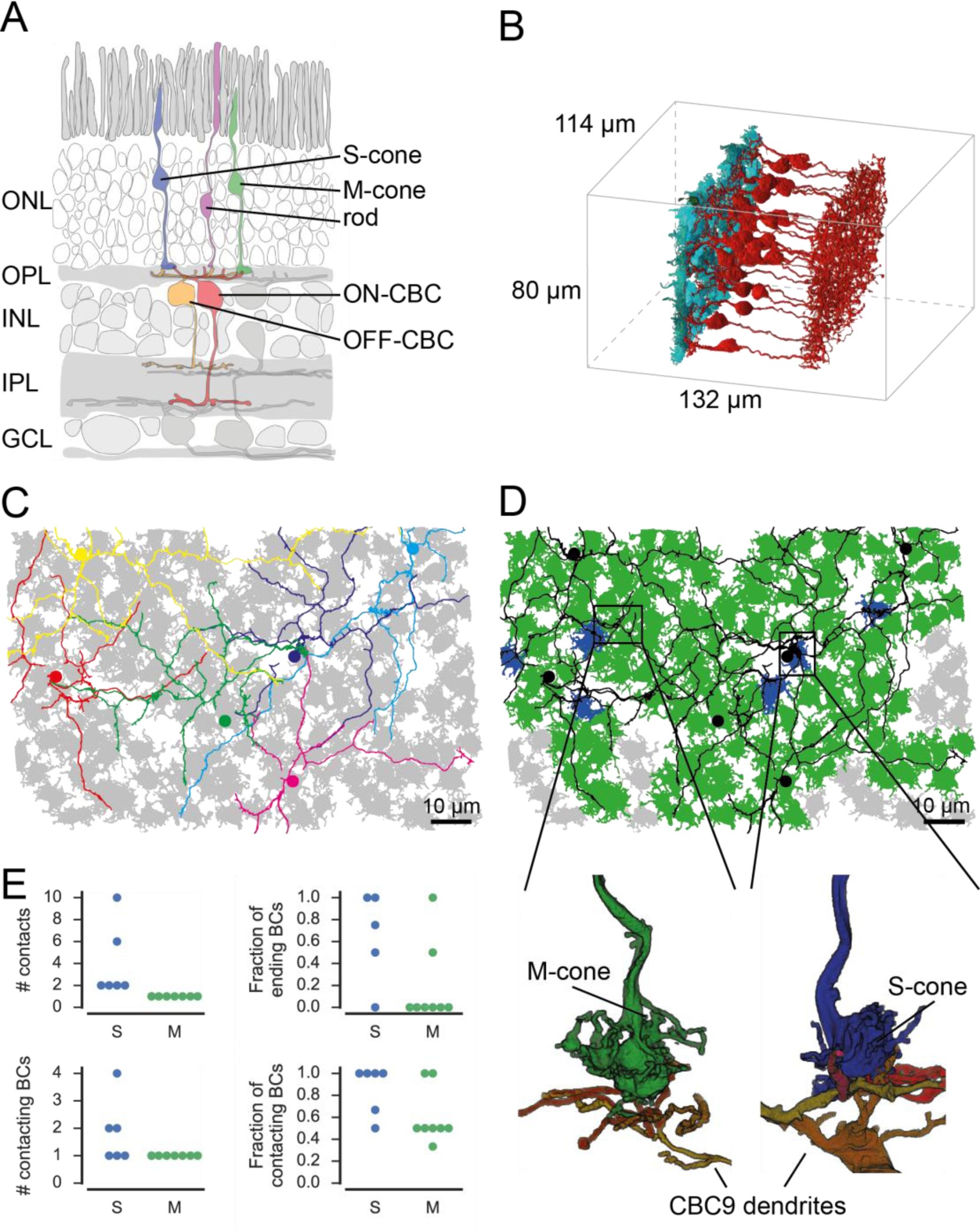
Identification of S-and M-cones. A. Scheme showing vertical section through the mouse retina. B. Volume-reconstructed cones and all CBC4 cells. C. Cone pedicles (grey) with CBC9s. BC soma localization is indicated by colored dots. D. Same as C, but with putative S-cones (blue) and M-cones (green) highlighted. Unidentified cones are shown in grey. Insets indicate the location of the examples shown below of cone pedicles contacted by CBC9 dendrites. E. Contact parameters used for S-cone identification. ONL, outer nuclear layer; OPL, outer plexiform layer; INL, inner nuclear layer; IPL, inner plexiform layer; GCL, ganglion cell layer.

To identify S-and M-cones we used the fact that type 9 cone bipolar cells selectively target S-cones (Figure 1C, D) (Mariani, 1984; Kouyama and Marshak, 1992; Haverkamp et al., 2005; Breuninger et al., 2011). We found 48 contacts of CBC9s and cones, involving 43 cones (Supplementary Figure 1A). We visually assessed all contacts and found that 29 of 79 these were in the periphery of the cone pedicle, where no synapses are expected (Supplementary Figure 1B) (Dowling and Boycott, 1966; Chun et al., 1996). This left 14 potential S-cones with invaginating contacts by at least one CBC9. We assumed that each S-82 cone is contacted by all CBC9 dendrites close to it and that those contacts occur mostly at the end of dendritic branches (Haverkamp et al., 2005). We excluded 8 potential S-cones according to these criteria (Figure 1E and Supplementary Figure 1C), resulting in 6 cones we 85 identified as S-cones (Figure 1D and Supplementary Figure 1D, see *Methods and Discussion*). This corresponds to a fraction of 4.8% S-cones (6/124 cones within the dendritic field of at least one CBC9), matching the 3-5% reported in previous studies (Röhlich et al., 1994; Haverkamp et al., 2005).

### Classification of photoreceptor-BC contacts

We next developed an automatic method to distinguish contacts likely corresponding to synaptic connections from false contacts. As the tissue in the dataset is stained to enhance cell-surface contrast in order to enable automatic reconstruction, it is not possible to distinguish between synaptic contacts based on explicit ultrastructural synaptic markers, such as vesicles, synaptic ribbons or postsynaptic densities (see also discussion in Helmstaedter et al., 2013). In contrast to the synaptic contacts in the inner plexiform layer studied by Helmstaedter et al. (Helmstaedter et al., 2013), the special morphology of synapses at cone pedicles still allowed us to classify the contacts (Haverkamp et al., 2000): The ribbon synapses of the cones are placed exclusively in the presynaptic area at the bottom of the cone pedicles. Here, ON-cone bipolar cells (ON-CBCs) make invaginating contacts, where the dendritic tips reach a few hundred nanometers into the presynaptic area of cone pedicles (Figure 2A) (Dowling and Boycott, 1966). In contrast, OFF-cone BCs (OFF-CBCs) make basal contacts in the same area (Figure 2B). These “true” contacts have to be distinguished from contacts in the periphery or at the (out)sides of the cone pedicle as well as contacts between dendrites and cone telodendria, which can happen, for instance as dendrites pass by (Figure 2C).

**Figure 2:**
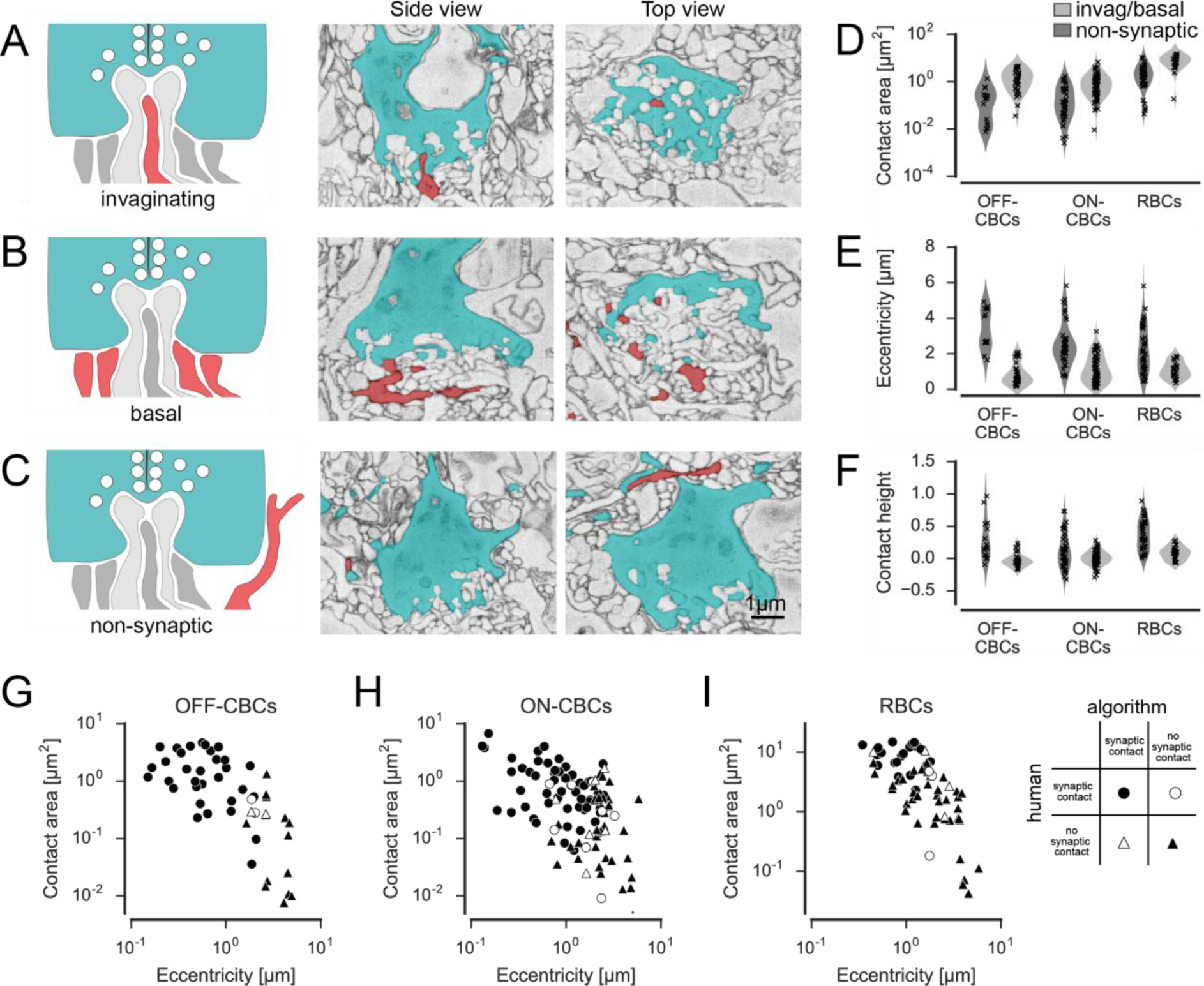
Classification of cone contacts. A. Invaginating ON-CBC contact. Schematic drawing (left), EM side view (center) and top view (right). Red and grey, BC dendrites; light grey, horizontal cell dendrites; cyan, cone pedicles. B. Basal/flat OFF-CBC contact as in A. C. Peripheral (non-synaptic) BC contact as in A. D.-E. Contact area (D), eccentricity (E), contact height (F) of invaginating/basal and non-synaptic contacts for OFF-/ON-CBCs and rod bipolar cells (RBCs). G.-I. Contact area versus eccentricity for OFF-CBC (G), ON-CBC (H) and RBC (I) contacts indicating correctly and incorrectly classified contacts.

In total, we found n=20,944 contacts in n=2,620 pairs of cones and BCs. We trained a support vector machine (SVM) classifier to distinguish whether or not an individual BC obtains input from a cone (as opposed to classifying each individual contact site, see *Methods*). To this end, we defined a set of seven features, such as contact area, eccentricity and contact height, which allowed distinguishing between potential synaptic contacts and “false” contacts (Figures 2D-F), and used a set of randomly selected manually labeled contacts (n=50 for OFF-CBCs, n=108 for ON-CBCs and n=67 for RBCs) as training data. We trained separate classifiers for ON-CBCs, OFF-CBCs and RBCs and found that the automatic classifiers could reliably distinguish between true and false contacts, with a success rate of ~90% (leave-one-out cross-validation accuracy, Figure 2 G-I).

### Contacts between cones and CBCs

We analyzed contacts between CBCs and S-and M-cones in the center of the EM stack where cones were covered by a complete set of all BC types. There was no difference in the number of CBCs contacted by S-and M-cones with 12.2 ± 1.5 CBCs (n=5 cones, mean ± SEM) for S-cones and 12.2 ± 0.4 CBCs (n =71 cones) for M-cones, respectively. Similarly, the total number of contact points per cone was almost identical for S-and M-cones with an average of 108 ± 24 per S-and 105 ± 5 per M-cone.

To study convergence patterns from cones onto individual CBCs in more detail, we analyzed the number of contacted S-and M-cones by an individual CBC of every type (Figure 3A and B). Most CBC types were contacted predominantly by M-cones, with an average of 2-6 cones contacting individual CBCs. One exception was the CBC9 that – by our definition of S-cones – received considerable S-cone input. We also detected a few contacts between CBC9s and M-cones; these are a consequence of our definition of S-cone and originate from those cones for which we found only single CBC9 contacts (see above, Figure 1 and *Discussion* for an alternative analysis).

**Figure 3:**
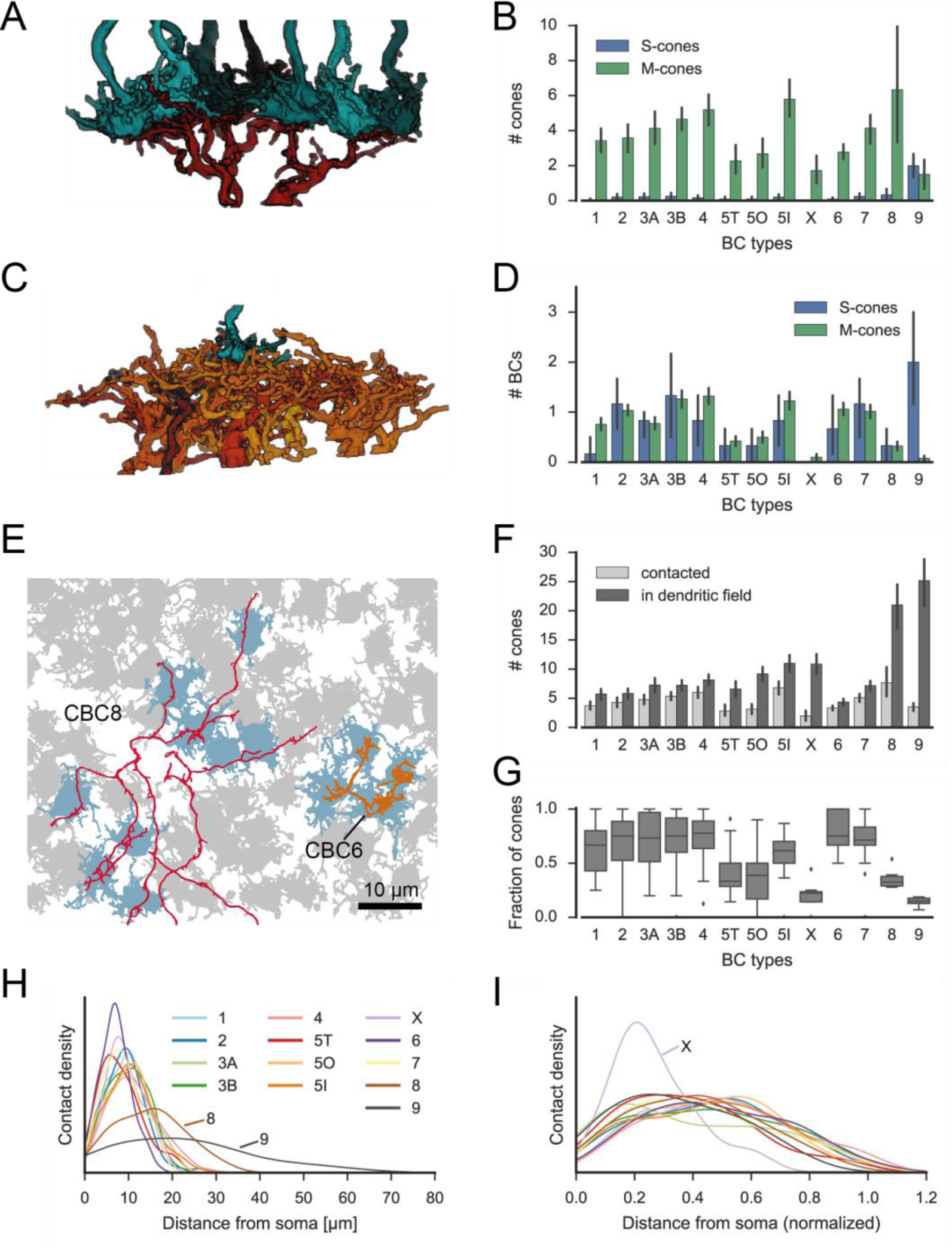
Quantification of cone-to-CBC contacts. A. Volume-reconstructed single BC dendrite (red) contacting numerous cone pedicles (cyan). B. Number of S-and M-cones contacted by different CBC types. C. Volume-reconstructed single cone (cyan) contacted by multiple BCs (orange/red). D. Number of CBCs per type contacted by individual S-and M-cones. E. Example cone array with CBC6 and CBC8 contacting cones. Grey, non-contacted cones; blue, contacted cones. F. Number of contacted cones and cones within dendritic field for different CBC types. G. Fraction of contacted cones/cones within dendritic field. H. Kernel density estimate of the distribution of contacted cones as function of distance from BC somata. I. Same as H. but distance normalized by dendritic field size. Bars in B,D,F indicate 95% CI.

To evaluate the divergent connectivity from S-and M-cones to CBCs, we studied how many individual BCs of each type were contacted by a single cone (Figure 3C). We found that each M-cone contacted on average a little less than one CBC1, while S-cones contacted almost no CBC1, consistent with previous reports (Breuninger et al., 2011). Conversely, we found that M-cones almost never contacted CBC9s, but S-cones contacted on average two. For all other CBC types, both cone types contacted them about equally (Figure 3D), with each cone making contact with at least one CBC2, 3B, 4, 5I, 6 and 7. In contrast, not every cone relayed its signals to every individual ON-CBC of types 5T, 5O, X and 8, as they were contacted by considerably less than one cone on average.

We next tested the hypothesis that CBCs other than type 1 and 9 unselectively contact all cones within their dendritic field (Wässle et al., 2009). To this end, we compared the number of contacted cones and the number of cones that are in reach of the BC dendrites (Figure 3E-G). OFF-CBCs (types 1-4) contacted on average 65-75% of the cones in their dendritic field, with very similar numbers across types (Figure 3G). In contrast, ON-CBCs showed greater diversity: The connectivity pattern of types 5I, 6 and 7 was similar to that observed in the OFF types (Figure 3G); these cells sampled from the majority of cones within their dendritic field (60-80%). CBC5T, 5O, X and 8, however, contacted less than half of the cones within their dendritic field (Figure 3G), with the lowest fraction contacted by CBCX (~20%). This result is not due to a systematic error in our contact classification: We manually checked volume-reconstructed dendritic trees of the respective types for completeness and frequently 4 found dendrites passing underneath a cone with a distance of 1-3 µm without contacting it(Supplementary Figure 2).

Finally, we studied the contact density along CBC dendrites (Figure 3H and I). To check for systematic variation independent of the absolute size of the CBC dendritic tree, we normalized the cone contact density for the dendritic field size of each CBC type (Figure 3I). Almost all CBC types received input at a very similar location relative to their soma, except for CBCX, which received the majority of inputs closer to the soma than all other types relative to its dendritic field size.

### The CBCX has few and atypical cone contacts

CBCX had an atypical connectivity pattern compared to other CBC types, we decided to study its connections in more detail. This BC type has only recently been identified by (Helmstaedter et al., 2013). It has a compact dendritic tree but a relatively wide axonal terminal system that stratifies narrowly at approximately the same depth as CBC5O and 5I do. Interestingly, CBCX seems to sample the cone input very sparsely, with input from only 2 cones on average, contacting only about 20% of the cones available in its dendritic field (Figure 3C, D and G). In fact, dendrites of CBCX oftentimes passed underneath cones or even stopped shortly before cone pedicles without making contacts at all (Figure 4A and B). It is unlikely that this resulted from incomplete skeletons for these BCs, as all skeletons were independently verified for this study and corrections were necessary (see Methods).

**Figure 4:**
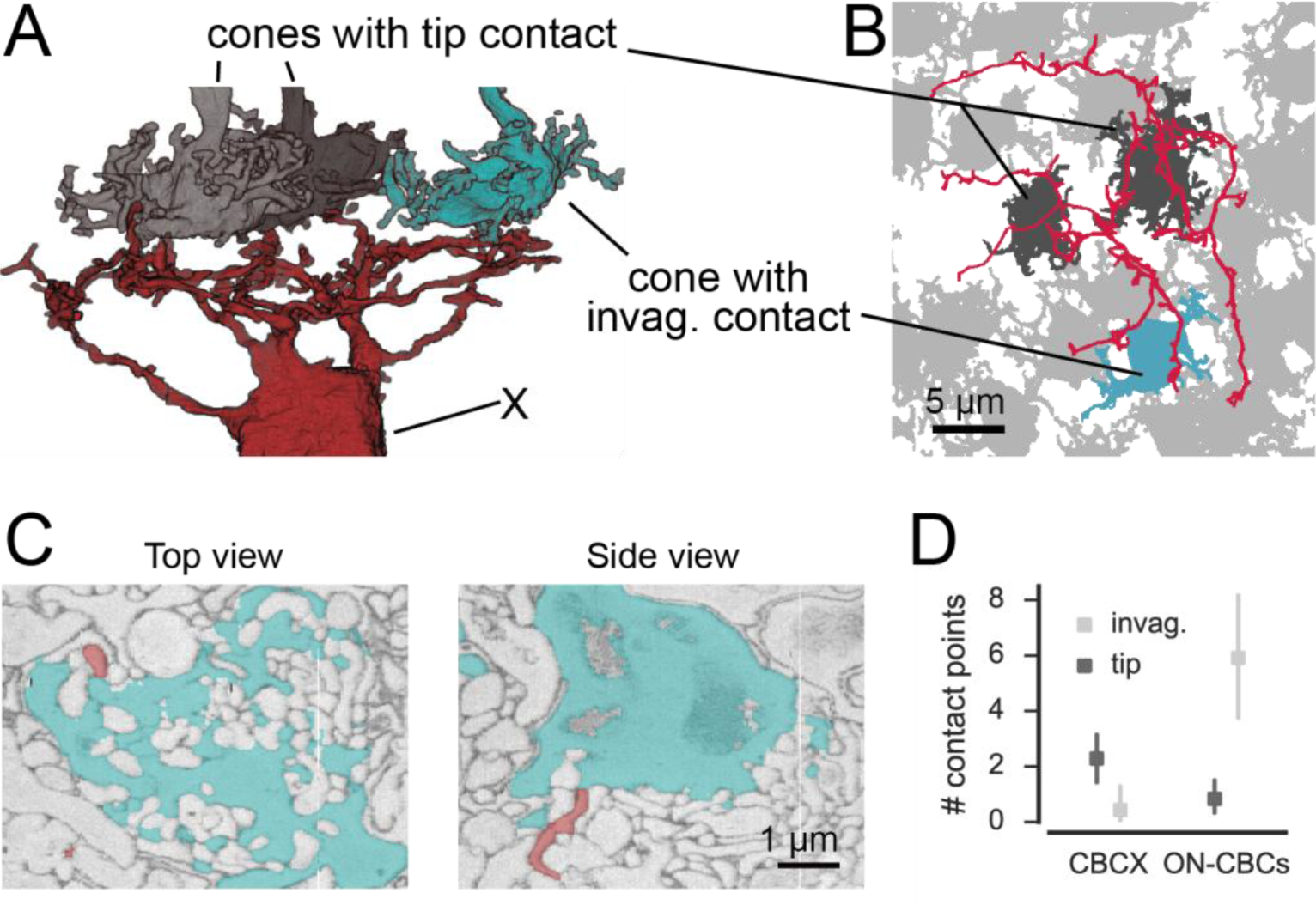
CBCX makes few and atypical cone contacts. A. Volume-reconstructed CBCX dendritic arbor (red) contacting few cone pedicles (cyan, invaginating contact; grey, tip contact). B. Example cone array as in A. with CBCX dendritic arbor contacting cones. Light grey, non-contacted cones; cyan, invaginating contacts, dark grey, tip contacts. C. EM image showing tip contact between CBCX (red) and cone pedicles (cyan), top view (left) and side view (right). D. Invaginating and tip contacts in CBCXs and other ON-CBCs. Bars in D. indicate 95% CI.

We re-examined all detected contacts between CBCXs and cones and found that very few of those were “classical” invaginating ON-CBC contacts (3 out of 19 contacts, n=7 cells, Figure 4B-D). The vast majority were “tip” contacts (16 out of 19 contacts, n=7 cells), which were similar to basal contacts made by OFF-CBC dendrites (Figure 4B-D). The available data was not conclusive with regards to the question whether these tip contacts of CBCX are smaller than those of OFF-CBCs (median area: 0.052 µm^2^ for n=22 CBCX contacts; 0.098 µm^2^ for n=23 OFF-CBC contacts, but p=0.17, Wilcoxon ranksum test).

In contrast to the CBCX, the other ON-CBC types made mostly invaginating contacts (71 out of 81 contacts, n=12 cells, 2 cells per BC type, Figure 4D), indicating a significant effect of cell type on contact type (GLM with Poisson output distribution, n=38, interaction: p<0.0001, see Methods). We checked if CBCX receive rod input instead but did not observe any rod contacts (see below). Thus, the CBCX appears to be an ON-CBC with both very sparse and atypical cone contacts similar to those made by OFF-CBCs. Still, based on the axonal stratification depth (Helmstaedter et al., 2013) and recent electrophysiological and functional recordings (Ichinose et al., 2014; Franke et al., 2016) this BC type is most likely an ON-CBC.

### RBCs make contacts with cones

We next analyzed the connectivity between photoreceptors and rod bipolar cells (RBCs) to test the hypothesis that RBCs may contact cones directly (Pang et al., 2010). In fact, RBCs did not only contact rod spherules but also cone pedicles (Figure 5A,B). These contacts were typical ON-CBC contacts with invaginating dendritic tips into the cone pedicles (Figure 5B). To quantify the cone-to-RBC connectivity in more detail, we counted the number of cones contacted by an individual RBC. While the vast majority (75%) contacted at least one cone, only 25% of all RBCs (n=141) did not contact any (Figure 5C). However, we did not find a preference of RBCs to connect S- or M-cones (Figure 5D). Conversely, 45% of cones contacted a single RBC, ~35% spread their signal to two to four RBCs, and only 20% of the cones did not make any contact with an RBC (Figure 5E). Our finding provides an anatomical basis to the physiologically postulated direct cone input into a subset of RBCs (Pang et al., 2010). Next, we evaluated whether RBCs contacting only rods or both cone(s) and rods represent two types of RBC, as hypothesized by Pang et al. (2010). However, the two groups of RBCs did not differ regarding their stratification depth (Supplementary Figure 3A), number of rod contacts (Supplementary Figure 3B) or potential connectivity to All amacrine cells (Supplementary Figure 3C), and did not form independent mosaics (Supplementary Figure 3D), arguing against two types of RBC.

**Figure 5:**
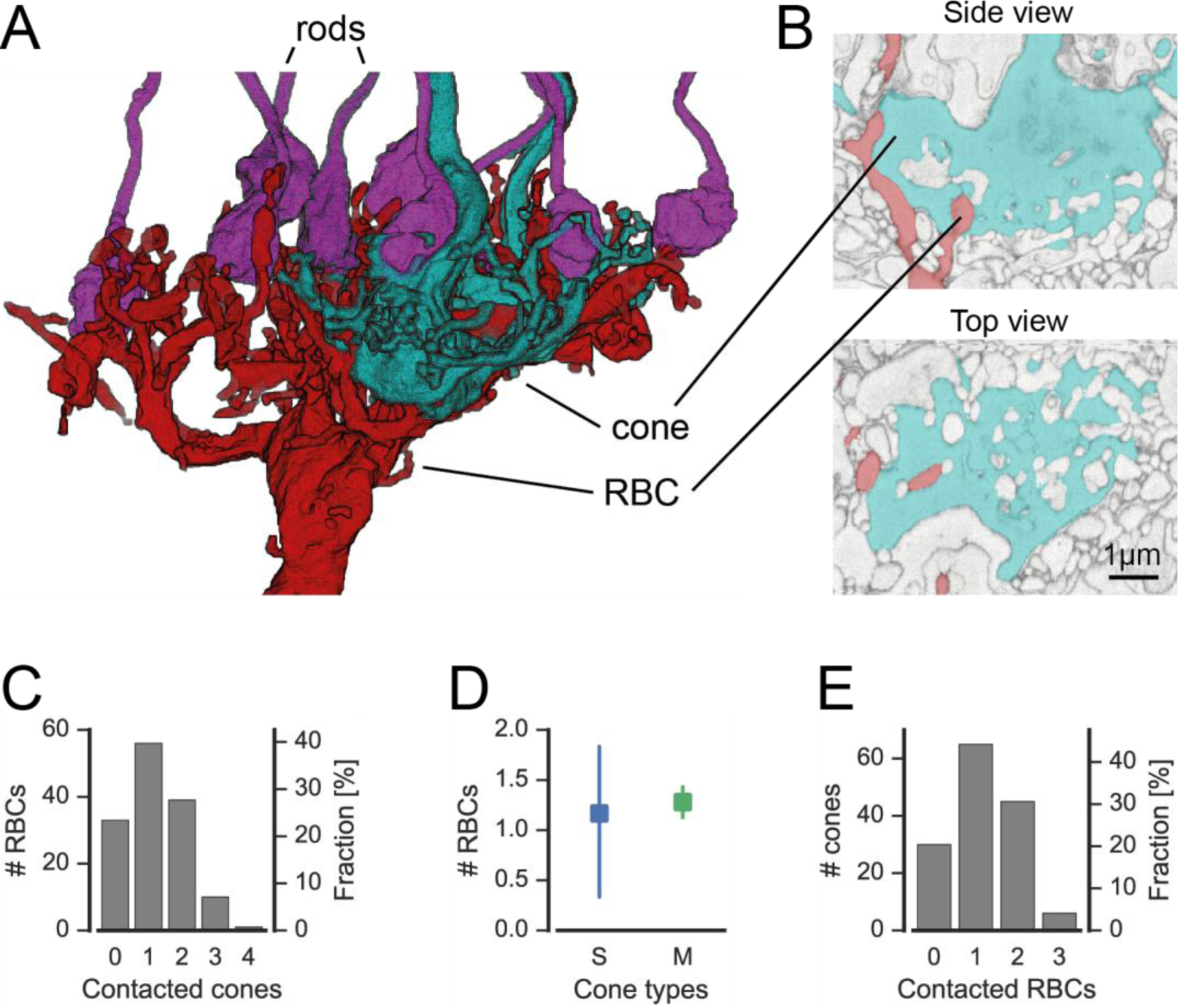
Cones contact rod bipolar cells. A. Volume-reconstructed RBC (red) contacting both rods (magenta) and cone pedicles (cyan). B. EM image showing invaginating contact between cone (cyan) and RBC (red), side view (top) and top view (bottom). C. Number of RBCs contacted by cones. D. Number of RBCs contacted by S- and M cones. E. Number of cones contacted by RBCs. Bars in D. indicate 95% CI.

### Quantification of rod to OFF-CBC contacts

Analogous to the analysis above, we skeletonized and volume rendered a complete set of over 2000 neighboring rod spherules (about 50% of the EM field, Figure 6A, Supplementary Figure 4) and identified rod-to-bipolar cell connections. In addition to the well-described invaginating rod-to-RBC connections (Figure 6B), we also found basal contacts between OFF-CBCs and rods close to the invaginating RBC dendrites (Figure 6C), as described earlier (Hack et al., 1999; Mataruga et al., 2007; Haverkamp et al., 2008; Tsukamoto and Omi,2014). We did not find any contacts between ON-CBCs and rods (in agreement with Tsukamoto and Omi, 2014; but see Tsukamoto et al., 2007).

**Figure 6:**
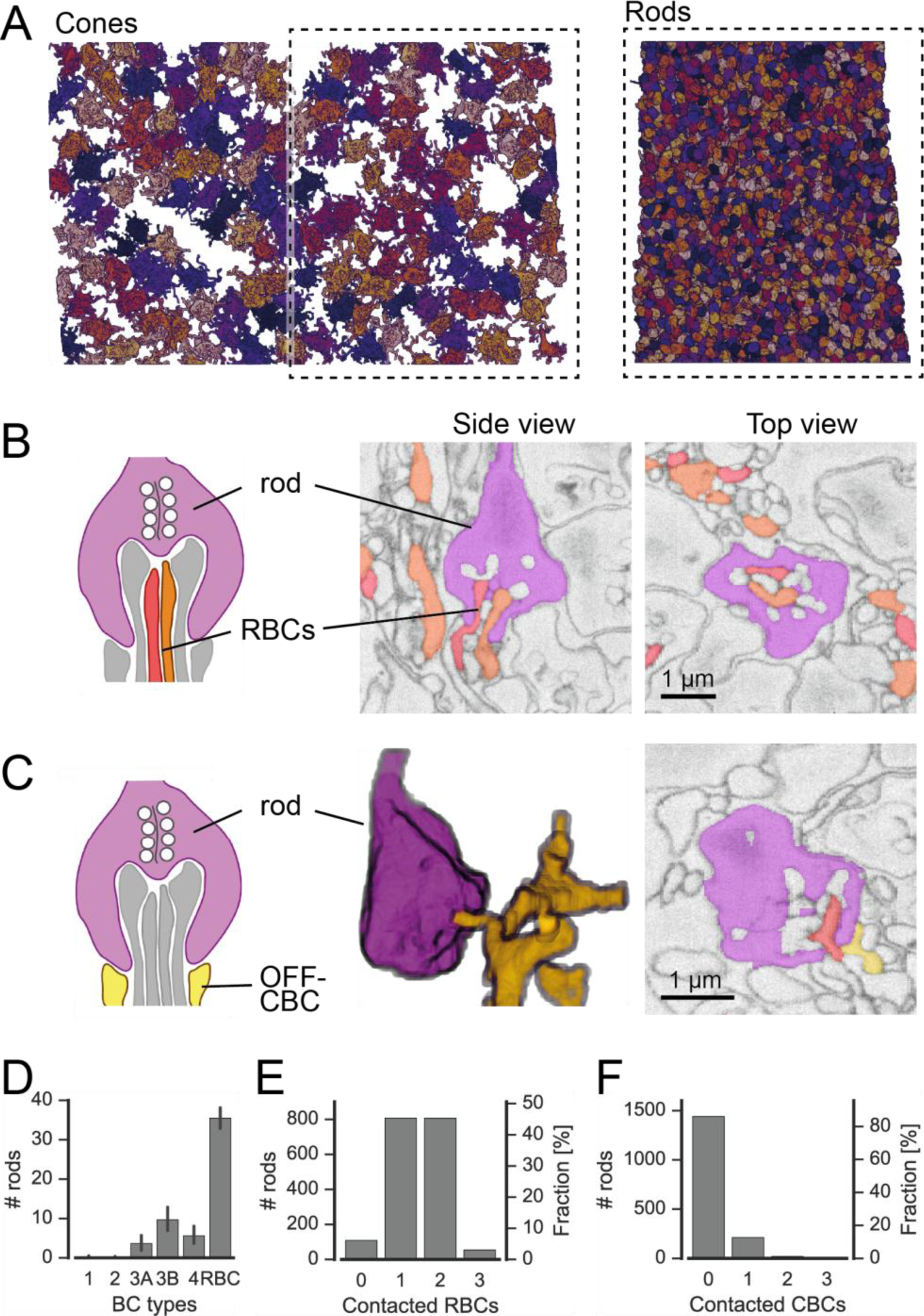
Rods contact RBCs and OFF-CBCs. A. Volume-reconstructed neighboring rod spherules (right) in one half of the field of the reconstructed cone pedicles (left). B. Rod spherule (magenta) with invaginating dendrites of two RBCs (orange, red). Schematic drawing (left), EM images side view (middle) and top view (right). C. Rod spherule (magenta) with basal contacts by OFF-CBCs (yellow). Schematic (left), volume-reconstructed vertical view (middle), EM image with top view (right). The latter also shows an invaginating RBC dendrite (red). D-F. Number of rods (and fraction) contacted by RBCs (D,E), and OFFCBC types (D, F). Bars in D. indicate 95% CI.

A single RBC contacted about 35 rods (Figure 6D), and a single rod contacted one or two RBCs, but very rarely no RBC or more than two (Figure 6E). In all cases with two invaginating dendrites, the dendrites belonged to two different RBCs (n=30 rods). The rods without RBC contacts were mainly located at the border of the reconstructed volume, where we could not recover all RBCs. The number of rods contacting OFF-CBCs was much lower: Whereas CBC1 and CBC2 did not receive considerable rod input, CBC3A, CBC3B and CBC4 were contacted by 5-10 rods, with CBC3B receiving the strongest rod input (Figure6D).

### Discussion

We analyzed an existing electron microscopy dataset (Helmstaedter et al., 2013) to quantify the connectivity between photoreceptors and bipolar cells. We found interesting violations of established principles of outer retinal connectivity: The newly discovered CBCX (Helmstaedter et al., 2013), likely an ON-CBC (Ichinose et al., 2014; Franke et al., 2016), had unexpectedly few and mostly atypical basal contacts to cones. While CBC types 5T, 5O from their dendritic field,and 8 also contacted fewer cones than expected from their dendritic field, they exhibited “standard” invaginating synapses. In addition, we provide anatomical evidence that rod and cone pathways are interconnected in both directions, showing frequent cone-RBC contacts.

### Does a ‘contact’ represent a synaptic connection?

Since the dataset we used was not labeled for synaptic structures, we used automatic classifiers based on structural and geometrical criteria to identify putative synaptic contacts between BCs and photoreceptors. These criteria allow unambiguous identification of synaptic sites for trained humans. For example, we used as a feature the proximity of the closest contact to the center of the cone pedicle region, where presynaptic ribbons have been reported at ultrastructural level (Dowling and Boycott, 1966; Chun et al., 1996). The overall accuracy of the classifiers evaluated with human annotated labels was high (~90%). Nevertheless, it is possible that a few contacts were misclassified. Manual quality control, however, revealed no systematic errors, indicating that all BC types should be affected similarly by any error in contact classification. For reference, all data including software for classifying and examining BC-cone contacts is available online. We believe that the “false contacts” are indeed due to dendrites of BCs passing by the cone pedicle and accidentally touching it; however, with the present dataset the existence of gap junctions at these contact points in at least some of the cases cannot be ruled out.

### Is there an effect of retinal location?

Unfortunately, the retinal location of the EM stack used here is unknown (Helmstaedter et al.,2013); it may originate from the ventral retina, where M-cones co-express S-opsin (Röhlich et al.,1994; Baden et al., 2013) However, “true” S-cones seem to be evenly distributed across the retina (Haverkamp et al., 2005), and hence CBC9 connectivity can be used for identification of S-cones independent of location. Nevertheless, it is possible that opsin co-expression in M-cones in the ventral retina influences the connectivity patterns between the M-cones and the remaining bipolar cell types.

### Alternative, more liberal S-cone classification

An alternative scheme for identifying S-cones would have been to classify all cones with invaginating contacts from CBC9 as S-cones, not only those with multiple, strong contacts. This would have resulted in a total of 14 S-cones out of 124 cones (Supplementary Figure 5A) or a fraction of 11.3%. Assuming a fraction of 3-5% S-cones (Haverkamp et al., 2005), this scenario is very unlikely (p=0.0037, binomial test, null hypothesis: 5% S-cones, n=124). We nevertheless ran the connectivity analysis with this set of S-cones (Supplementary Figure 5B). In this analysis, CBC9 was the only color specific BC type whereas all other BC types including CBC1 contacted both S-and M-cones without preferences (Supplementary Figure 5C), contradicting previous physiological findings (Breuninger et al., 2011).

### Sparse contacts between some ON CBC types and cones

We found that ON-CBCs 5T, 5O, X and 8 contact less cones than expected from the size of 4 their dendritic field. We observed that many of their dendrites passed by the cone pedicles with a distance of 1-3 µm or even ended under a cone pedicle without contacting it (Supplementary Figure 2). This is in agreement with a recent study reporting that CBC8s do not contact all cones within their dendritic field (Dunn and Wong, 2012), but in contrast to earlier studies that concluded that diffuse BCs receive input from all cones within their dendritic field (Boycott and Wässle, 1991; Wässle et al., 2009). However, a crucial difference with the earlier studies and our study is the spatial resolution: Whereas conventional confocal microscopy can resolve depth with a resolution of several hundreds of nanometers, the EM dataset we used has a resolution of 25 nm, allowing us to more accurately assess whether pre- and postsynaptic structures are in contact with each other.

It is also possible that some ON-CBC types make additional ‘diffusion-based’ synaptic contacts very similar to what has been described for OFF-CBCs (DeVries et al., 2006), for diffusion between cones (Szmajda and DeVries, 2011) or volume-transmitting neurogliaform 277 cells in cortex (Jiang et al., 2015). Thus, a lack of a membrane-to-membrane contact between a cone and a BC dendrite may not necessarily indicate the absence of synaptic signaling.

### CBCX makes atypical contacts with cones

As shown above, the CBCX makes even fewer contacts with cones: On average, they contact only about two cones, representing a fraction of only 20% of the cones within the area of their dendrites. Interestingly, this finding is in agreement with a recent report of single-cell RNA-seq experiments that CBCXs show clearly lower expression levels for metabotropic glutamate receptor *mGluR6* (*grm6*) – the hallmark of ON-BCs – compared to other ON-CBC types (J. Sanes, personal communication). This behavior is reminiscent of the giant CBC in macaque retina (Joo et al., 2011). Like the CBCX in mouse retina, it has a very large and sparsely branched dendritic tree and a relatively large axonal arbor that stratifies in the middle of the IPL and contacts only about 50% of the cones in its dendritic field.

In contrast to all other ON-CBCs, we found that the vast majority of CBCX contacts were not invaginating but tip, rather resembling basal OFF-CBC contacts. It is unclear whether or not these tip contacts are indeed functional synaptic sites. This is not the first finding to challenge the traditional view that ON-CBCs form only invaginating and OFF-CBCs only basal synaptic contacts. In the primate fovea, diffuse ON-CBCs (DBs) form basal contacts with foveal cones since almost all invaginating sites host midget bipolar cell dendrites (Calkins et al., 1996). This space limitation is less evident in mid-peripheral primate retina. At 3-4 mm eccentricity, diffuse ON-CBCs receive 10% (DB5) to 40% (DB4 and DB6) of their cone input through basal synapses (Hopkins and Boycott, 1996). As CBCX also expresses an AMPA-type basal contacts. However, direct glutamate receptor (*gria2;* J. Sanes, personal communication), it is possible that they receive ON input via invaginating and OFF input by flat/basal contacts. However, direct functional evidence that the CBCX can have an OFF signal component is lacking so far (Ichinose et al., 2014; Franke et al., 2016).

Interestingly, also CBCX contacts in the IPL also appear to be distinct from those of other BC types: First, the majority of cells contacted by CBCX in the IPL are amacrine cells rather than ganglion cells (Helmstaedter et al., 2013). Second, they form sparse contacts relative to their axon terminal size with comparatively few cells. Thus, the CBCX seems to be an exception, an unusual BC type in many respects in addition to its sparse and atypical connectivity properties in the OPL, reminiscent of the recently described dendrite-less bipolar cell(Della Santina et al., 2016).

### RBCs may form an additional photopic ON channel

We found that cones connect to 75% of the RBCs; in many cases, one cone contacted multiple RBCs. In turn, 35% of the RBCs received converging input from several cones. This massive cone input via invaginating synapses to RBCs suggests a prominent use of the primary rod pathway (Bloomfield and Dacheux, 2001) during photopic conditions. Consistent with our findings, it has been reported before that RBCs can be activated under photopic light conditions (Chen et al., 2014; Tikidji-Hamburyan et al., 2015; Franke et al., 2016), even when rods are “traditionally” expected to be fully saturated, but the functional significance of photopic RBC activity is not clear. RBCs could indirectly inhibit OFF-CBCs via All amacrine cells; as they will likely not activate ON-CBCs since the AII-ON-CBC gap junctions are believed to be closed under these conditions (Bloomfield et al., 1997); this suggests that RBCs contribute to crossover inhibition (Molnar and Werblin, 2007).

Based on the physiological finding that only a subset of RBCs receive input from cones, Pang et al. (2010) suggested that there may be two distinct RBC types, with the rod-only one having axon terminals ending closer to the ganglion cell layer. Our data does not provide evidence for two RBC types based on the connectivity in the outer retina (see Supplementary Figure 3). This agrees well with recent findings from single-cell RNA-seq experiments, where all RBCs fell into a single genetic cluster with little heterogeneity (J. Sanes, personal communication).

### OFF CBC types contact different numbers of rods

We quantified the number of rods contacting the five OFF-CBC types. Whereas CBC1 and 2 received almost no rod input, we observed flat/basal contacts between rods and types CBC3A, 3B and 4, providing a quantitative confirmation of this finding (Mataruga et al., 2007; Haverkamp et al., 2008; Tsukamoto and Omi, 2014). CBC3A and 4 received input from ~5 rods in addition to the ~5 cones contacted by them. CBC3B sampled from the same number of cones but was contacted by about twice as many rods. Thus, rods provide considerable inputs to OFF-CBCs, possibly representing a distinct scotopic OFF channel complementing the scotopic ON channel via RBCs. Interestingly, the morphologically very similar CBC3A and 3B may obtain their (functional) differences not only from the expression of different ionotropic glutamate receptors (Puller et al., 2013) but also from their connectivity with rods.

### Conclusion

Here, we performed a systematic quantitative analysis of the photoreceptor-to-bipolar cell synapse. We showed that there are exceptions to several established principles of outer retinal connectivity; In particular, we found several ON-BC types that contacted only a relatively small fraction of the cones in their dendritic field. We also, we find that rod and cone pathways already interact strongly in the outer plexiform layer. Whether these are general features of mammalian retinas or whether these exceptions are evolutionary specializations unique to the mouse remains to be seen.

## Meterials and methods

### Dataset and preprocessing

We used the SBEM dataset e2006 published by (Helmstaedter et al., 2013) for our analysis. The dataset has a voxel resolution of 16.5 × 16.5 × 25 nm with dimensions 114 µm × 80 µm ×132 µm. We performed volume segmentation of the outer plexiform layer (OPL) using the algorithms of (Helmstaedter et al., 2013). The preprocessing of the data consisted of three steps: (i) Segmentation of the image stack, (ii) merging of the segmented regions and (iii) collection of regions into cell volumes based on traced skeletons.

We modified the segmentation algorithm to prevent merging of two segments if the total volume was above a threshold (>50,000 voxels), as sometimes the volumes of two cone pedicles could not be separated with the original algorithm. Although this modification resulted in overall smaller segments, these were collected and correctly assigned to cells based on the skeletons in the last step of the preprocessing.

We identified 163 cone pedicles and created skeletons spanning their volume using the software KNOSSOS ((Helmstaedter et al., 2012), www.knossostool.org/. We typically traced the center of the cone pedicle coarsely and added the individual telodendria for detailed reconstruction. In addition, we traced 2,177 rod spherules covering half of the dataset (Figure 6). For our analysis, we used all photoreceptors for which at least 50% of the volume had been reconstructed (resulting in 147 cones and 1,799 rods). We used the BC skeletons published by Helmstaedter et al. (2013), with the following exceptions: We completed the dendritic trees of three XBCs (CBCXs), which were incompletely traced in the original dataset. In addition, we discarded three BCs originally classified as RBCs because they were lacking rod contacts as well as the large axonal boutons typical for RBCs (Supp. Fig. 6 A-C),and one BC classified as a CBC9 because its dendritic field was mostly outside of the data stack (Supp. Fig. 6D).

Next, we used the algorithm by (Helmstaedter et al., 2013) to detect and calculate the position and area of 20,944 contact points between cone pedicles and BC dendrites and 7,993 contact points between rod spherules and BC dendrites. To simplify the later visual inspection of contacts, we used the reconstructed cell volumes to generate colored overlays for the raw data to highlight the different cells in KNOSSOS.

### Identification of S-cones

We detected 169 contacts in 51 pairs of CBC9s and cones. Upon manual inspection, we found a total of 32 invaginating (potentially synaptic) contacts between 6 CBC9s and 14 cone pedicles.

Based on immunocytochemistry, it has been shown that S-cones are contacted by all CBC9 within reach and that CBC9 contacts to S-cones are mostly at the tips of the dendrites (Haverkamp et al., 2005). For all 14 contacted cones, we analyzed the number of invaginating CBC9 contacts, the number of contacting CBC9s, the fraction of CBC9 with dendrites close to the cone that make contact and whether the dendrites end at the cone or continue beyond it (Figure 1E). Based on these criteria, we classified 6 out of these 14 cones as S-cones. In addition to our main analysis, we present an alternative analysis that considers the case if all 14 cones were counted as S-cones (Supplementary Figure 5).

### CBC5 classification

CBC5s were classified initially based on their connectivity to ganglion cells and amacrine cells into types 5A and 5R, where 5R was a group containing multiple types (Helmstaedter et al.,2013). In addition, some CBC5s could not be classified due to a lack of axonal overlap with the reconstructed ganglion cells of the types used for classification. Considering the separate coverage factors for dendritic and axonal overlap of all CBC5s together (OPL: 3.14, IPL: 2.89), dividing them into three subtypes is conceivable considering the numbers for other CBC types (Supplementary Table 3). This has already been suggested by (Greene et al.,2016), who divide CBC5s into three subtypes based on axonal density profiles (using a different EM dataset that includes only the inner retina).
We followed the classification approach suggested by Greene, Kim and coworkers (Greene et al. 2016): First, we calculated the densities of both ON-and OFF-starburst amacrine cells (SACs) dendrites along the optical axis. We fitted the peak of these profiles with a surface using bivariate B-splines of third order. Next, we corrected the density profiles of CBC5 axonal trees by mapping the SAC surfaces to parallel planes. We them applied principal component analysis (Supplementary Figure 7A) to obtain a first clustering into three groups by fitting a Gaussian mixture model (GMM) (Bishop, 2006) with three components onto the first three principal components of the axon density profiles. The resulting density profiles of the three clusters matches those found by (Greene et al., 2016) (Supplementary Figure 7B). As we noted a few violations of the postulated tiling of the retina by each type (Seung and Sümbül, 2014), we implemented a heuristic to shift cells to a different cluster or swap pairs of cells optimizing a cost function including both overlap in IPL and OPL as well as the GMM clustering (Supplementary Figure 7C):

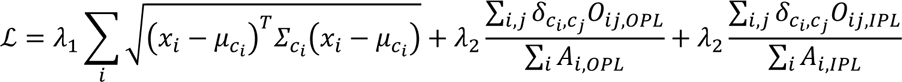

With *x_i_* the parameter vector of cell *i, c_i_*, the mixture component cell *i* is assigned to, μ_c_ the mean of the mixture component *c, σ_c_* the covariance matrix of the mixture component *c, g_ij_* the Kronecker delta, *A_i,OPL/IPL_* the area of the dendritic field/axonal tree of cell *i* and *O_ij,OPL/IPL_* the overlap of cell *i* and *j* in the OPL/IPL. The overlap of two cells is calculated as the intersection of the convex hull of the dendritic fields/axonal trees.

### Automatic contact classification

To distinguish potential synaptic contacts between photoreceptors and BCs from accidental contacts, we developed an automatic classification procedure exploiting the stereotypical anatomy of cone-BC synapses (triads, (Dowling and Boycott, 1966)). First, we grouped all contacts for a specific cone-BC pair, in the following referred to as a contact-set. We obtained a training data set by randomly selecting 10 contact-sets per CBC type and 50 RBC-cone contact-sets. We excluded CBCX from the training data because of their atypical contacts. To increase classifier performance we added 17 additional RBC-cone contact-sets 12 manually classified as invaginating contacts as well as all 48 CBC9-cone contact-sets classified for the S-cone identification. For those contact-sets, we visually inspected each individual contact point in the raw data combined with volume segmentation overlay using KNOSSOS. Then we classified it either as a central basal contact (potentially synaptic) or peripheral contact (e.g. at the side of a cone or contact with telodendria, likely non-synaptic)for OFF CBCs or as invaginating contact vs. peripheral contact for ON CBCs and RBCs. Next, we extracted a set of seven parameters for each contact (see Supplementary Figure 8):

- Contact area: The total contact area aggregated over all contact points between a BC and a cone
- Eccentricity: The distance between the cone center and the closest contact point in the plane perpendicular to the optical axis
- Contact height: The distance of the contact point with minimal eccentricity from the bottom of the cone pedicle (measured along the optical axis, normalized by the height of the cone pedicle).
- Distance to branch point: Minimal distance between a contact point and the closest branch point, measured along the dendrite of the cone pedicle).
- Distance to tip: Minimal distance between a contact point and the closest dendritic tip. A large distance occurs for example for a contact between a passing dendrite and a cone.
- Smallest angle between the dendrite and the optical axis at a contact point
- Number of contact points between cone and BC

Based on those parameters we trained a support vector machine classifier with radial basis functions (C-SVM) for each OFF-CBC, ON-CBC and RBC cone contact using the Python package *scikit-learn*. Optimal parameters were determined using leave-one-out cross validation (see Supplementary Table 1 for scores and error rates).

### Analysis of rod contacts

As the reconstructed rod spherules cover only half of the EM dataset, we restricted the analysis to bipolar cells with their soma position inside this area. To automatically classify the contacts to rods, we followed a similar scheme as for the cones. Again, we grouped the contacts for each pair of BC and rod spherule. As training data, we selected all putative contact sites with CBC1s (n=5) and CBC2s (n=32), 20 random contacts to CBC types 3A, 3B and 4 as well as 100 random contacts to RBCs. Again, we classified these contacts by visual inspection in KNOSSOS using the raw data with colored segmentation overlay. In addition, we manually inspected all 132 contact points between rod spherules and ON-CBCs, but could not identify a single potential synaptic contact. We trained SVM classifiers for contacts between rods and RBCs/OFF-CBCs using the same parameters as for the contacts to cones. As synaptic contacts between OFF-CBCs and rod spherules are basal contacts situated close to the invaginating RBC contacts, we added the minimum distance to the next (synaptic) RBC contact as additional classification parameter for OFF-CBCs. As a consequence, we restricted the analysis of OFF-CBC-to-rod contacts to those rods were cone. 13 RBC contacts could be identified (n=1,685). See Supplementary Table 2 for scores and error rates from the leave one out cross validation.

### Statistics

Error bars in all plots are 95% confidence intervals (CI) calculated as percentiles of the bootstrap distribution obtained via case resampling. In Figure 4D, we used a generalized linear mixed model with Poisson output distribution and fixed effects contact type and cell type and random effect cell identity (R package *lme4*). The model yielded a significant intercept (z=8.72, p<.0001), a significant main effect of cell type (z=-4,11, p<.0001), a significant main effect of contact type (z=-5.80, p<.0001) and a significant interaction cell x contact type (z=5.09, p<.0001).

### Data and code availability

All BC and PR skeletons including updated type annotations as well as connectivity data will be available as Supplementary Material (S Data 1 and 2). Jupyter notebooks for reproducing analysis and main figures will be available online.

**Figure 7:**
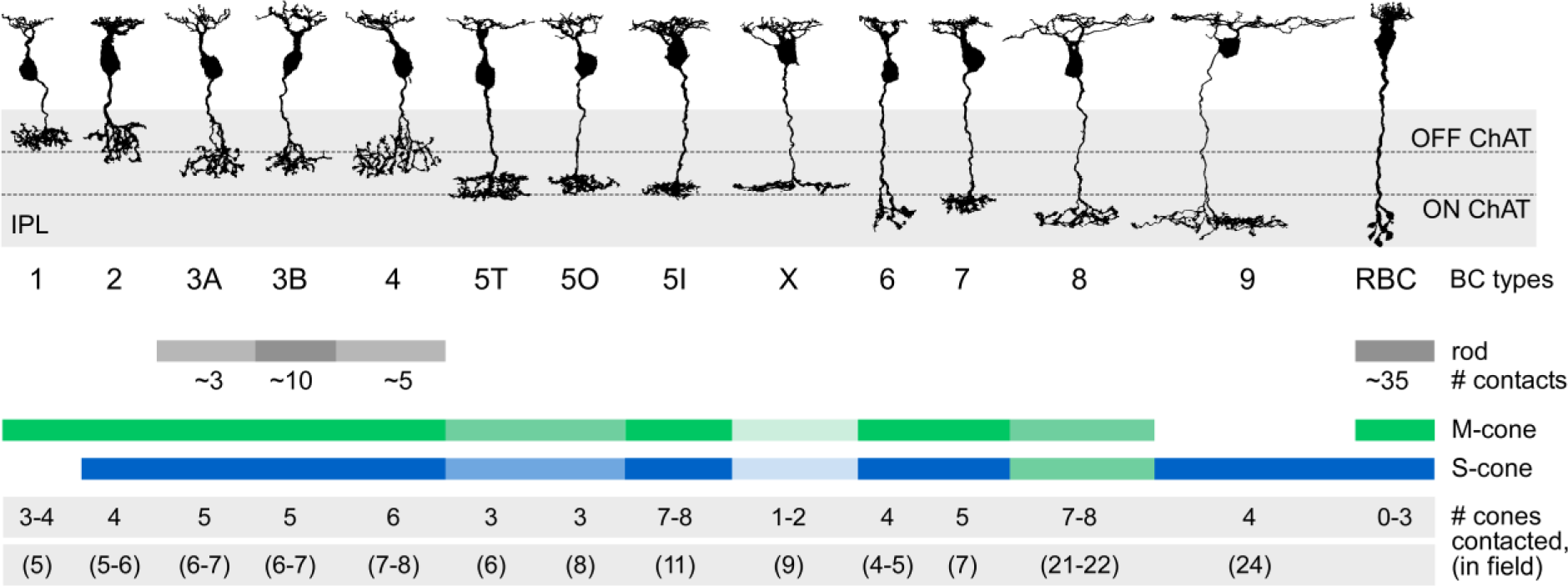
Connectivity between cone and rod photoreceptors and bipolar cells in the mouse retina. Representative examples of bipolar cell types in the mouse retina are shown. The number of cones in the dendritic field number and contacted photoreceptors are given for each type.

**Table 1:** Cross validation results of BC-to-cone contact classification

|  | False positive | False negative | Total score |
| --- | --- | --- | --- |
| OFF-CBCs | 12.5 % | 5.9 % | 0.92 |
| ON-CBCs | 14.0 % | 12.3 % | 0.87 |
| RBCs | 9.3 % | 12.5 % | 0.90 |

**Table 2:** Cross validation results of BC-to-rod contact classification

|  | False positive | False negative | Total score |
| --- | --- | --- | --- |
| OFF-CBCs | 18.3 % | 22.5 % | 0.8 |
| RBCs | 14.3 % | 2.6 % | 0.95 |

**Table 3:** OPL hull area: Average area of convex hull of dendritic field in OPL per cell type [µm^2^], mean ± SEM; OPL cov.: coverage factor derived from convex hulls by computing the sum of convex hull areas divided by area of the union of convex hulls; OPL cov. cones: coverage factor computed from cones by computing the sum of the number of cones in the dendritic field of each cell divided by the number of cones in the joint dendritic field; Wässle: coverage values from Wässle et al. 2009 computed by the same method as OPL cov. cones; IPL hull area: Average area of convex hull of the axonal field in IPL per cell type [µm^2^], mean ± SD; IPL cov: analogous to OPL cov.

| Type | n | OPL hull area<br>[$\mu\text{m}^2$] | OPL<br>cov. | OPL cov.<br>cones | Wässle | IPL hull area | IPL<br>cov. |
| --- | --- | --- | --- | --- | --- | --- | --- |
| CBC1 | 26 | 175±16 | 1.17 | 1.48 | 1.48 | 376±16 | 1.52 |
| CBC2 | 34 | 204±19 | 1.18 | 1.55 | 1.5 | 353±23 | 1.52 |
| CBC3A | 22 | 273±28 | 1.17 | 1.37 | 1.25 | 308±28 | 1.21 |
| CBC3B | 32 | 292±19 | 1.41 | 1.90 | 1.55 | 224±9 | 1.24 |
| CBC4 | 30 | 302±20 | 1.32 | 1.86 | 1.6 | 274±23 | 1.33 |
| CBC5T | 22 | 256±30 | 1.13 | 1.30 | - | 402±25 | 1.28 |
| CBC5O | 22 | 380±41 | 1.35 | 1.60 | - | 359±23 | 1.17 |
| CBC5I | 25 | 459±30 | 1.55 | 1.95 | - | 276±14 | 1.22 |
| CBCX | 7 | 433±34 | 1.02 | 1.12 | - | 899±126 | 1.12 |
| CBC6 | 45 | 125±11 | 1.14 | 1.58 | - | 165±11 | 1.17 |
| CBC7 | 29 | 254±18 | 1.22 | 1.65 | 1.3 | 274±11 | 1.16 |
| CBC8 | 6 | 1249±144 | 1.14 | 1.21 | - | 699±55 | 1.02 |
| CBC9 | 6 | 2223±227 | 1.84 | 1.45 | - | 1605±335 | 1.43 |
| RBC | 141 | 128±3 | 2.17 | 4.37 | - | 65±3 | 1.40 |

## Acknowledgements

### Acknowledgements

We thank M. Helmstaedter and coworkers (2013) for making their data available. This work was funded by the DFG (EXC 307 and BE 5601/1-1) and the BMBF through the BCCN Tübingen (FKZ 01GQ1002) and the Bernstein Award to PB (FKZ 01GQ1601).

## Author contributions

TS, SH, TE and PB designed the study; CB analyzed the data; TS and CB performed anatomical tracing; TE and PB supervised the study; all authors contributed to writing the manuscript.

---

**Supplementary Figure 1:**
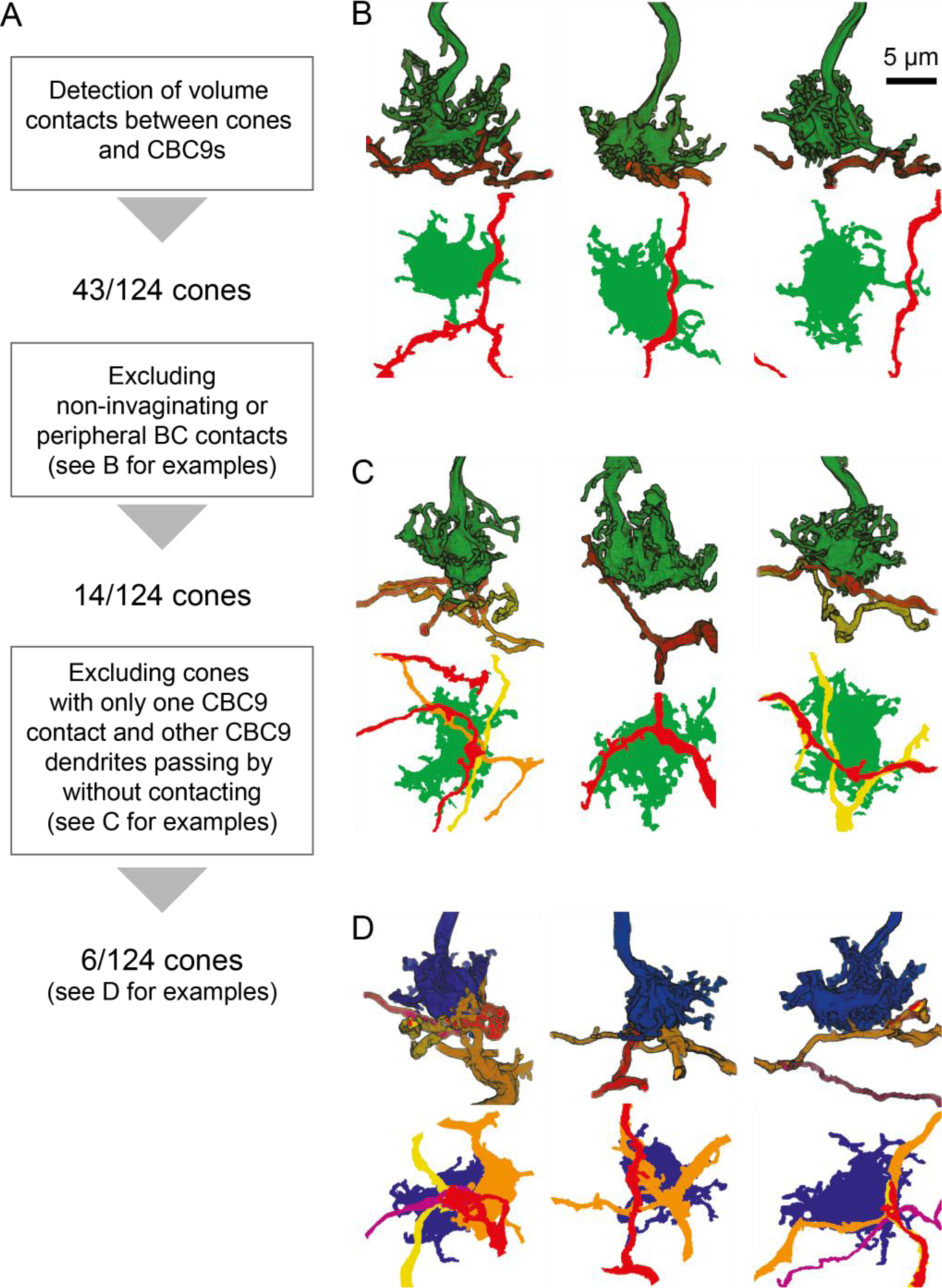
A. Diagram showing workflow for identification of S-and M-cones using connectivity with CBC9 cells. B.-D. Side view and horizontal projection of representative examples of cone pedicles (green, M-cone; blue, S-cone) with CBC9 dendrites (yellow, orange, red) with non-invaginating but peripheral contacts (B), with only one CBC9 contact and other CBC9 dendrites passing by (C) and ‘true’ S-cones (D).

**Supplementary Figure 2:**
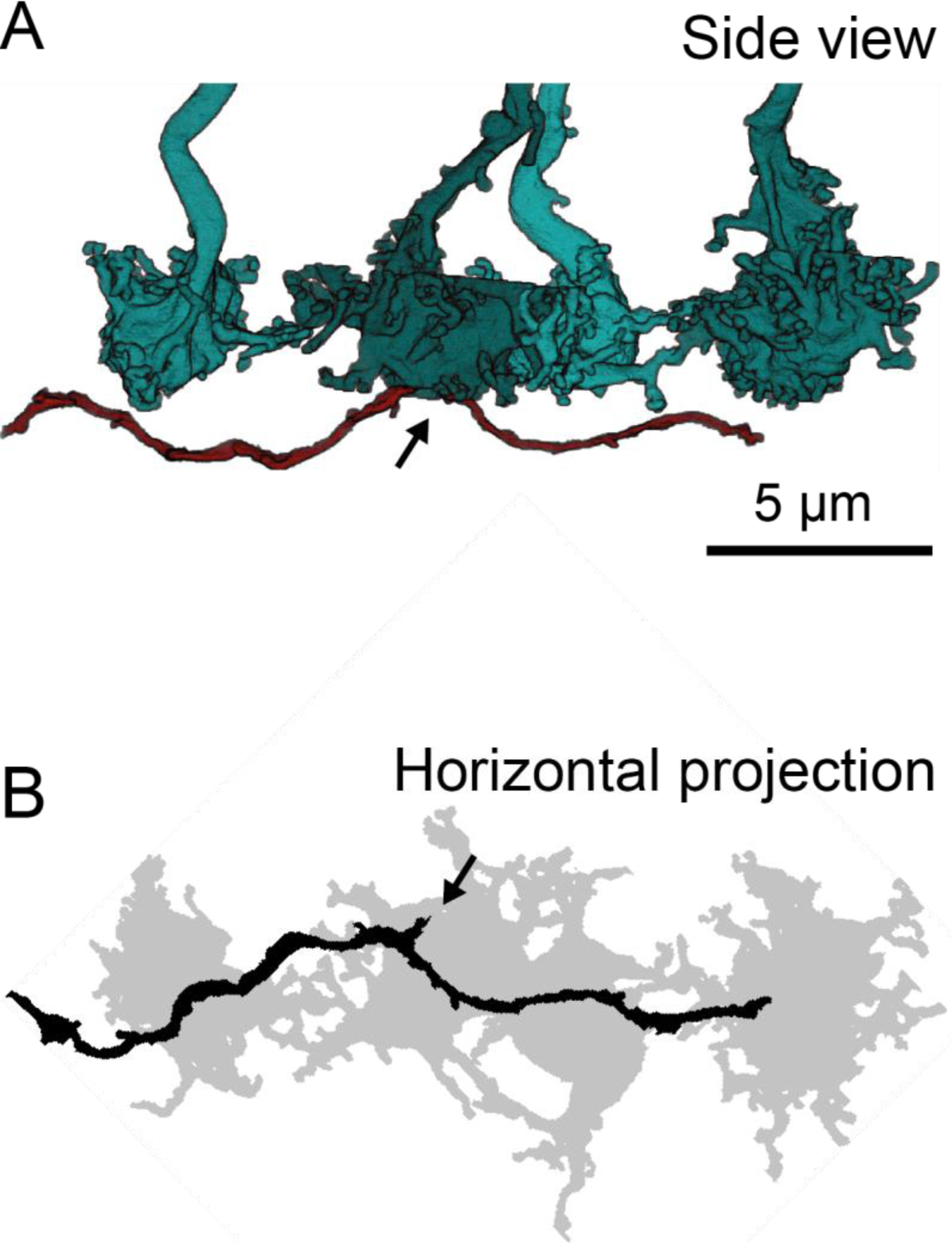
A. Side view of four volume-reconstructed cone pedicle (cyan) and CBC8 dendrite (red). B. Horizontal projection of the neurite structures shown in A. Arrow indicates invaginating ON-CBC contact.

**Supplementary Figure 3:**
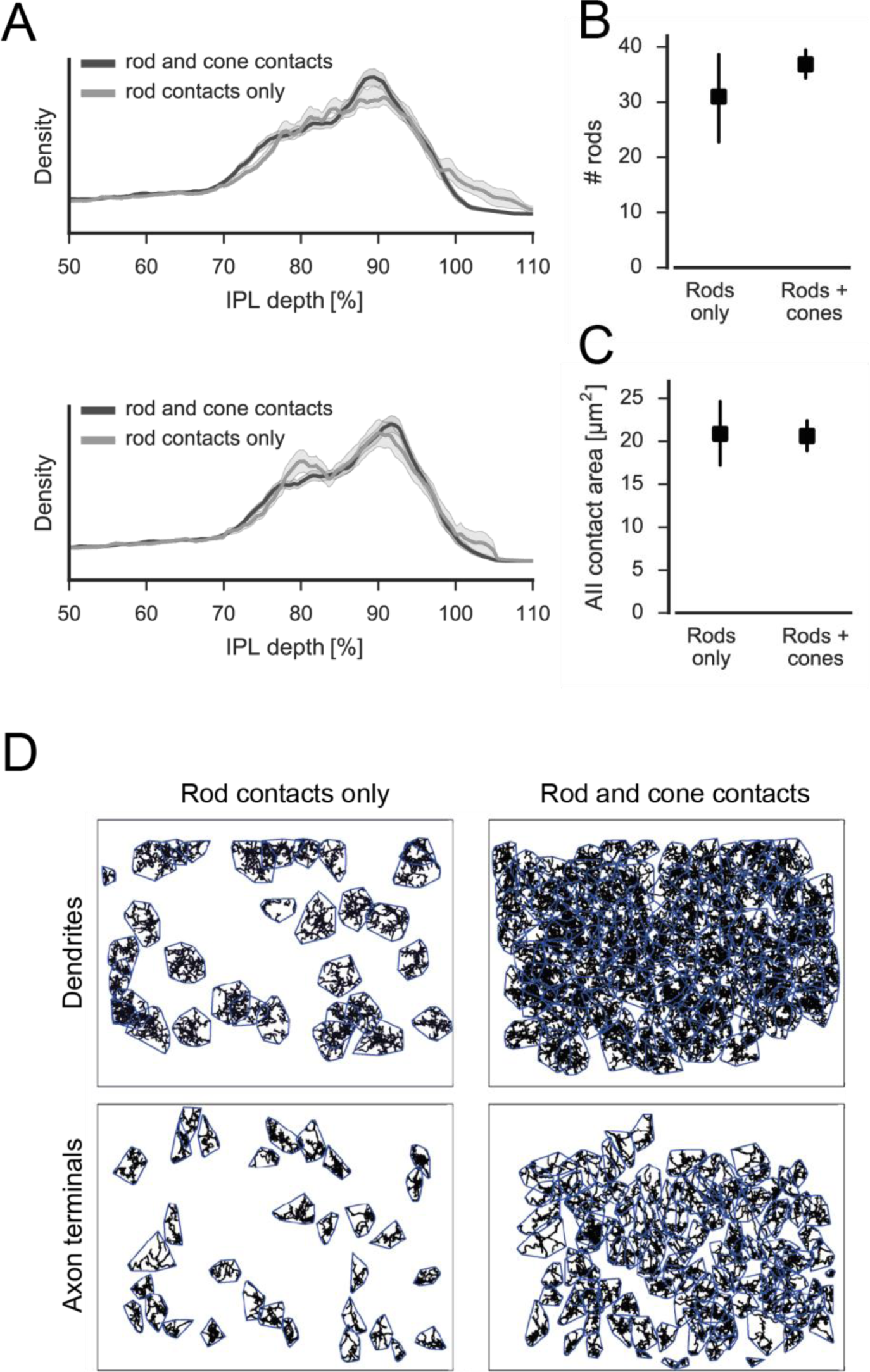
A. Relative density of RBC spherules in the IPL using both dendritic ON and OFF starburst amacrine cell (SAC) bands (top) and only the dendritic ON SAC band (bottom) for depth correction (shading: SEM). B. Number of rods contacted by RBCs contacting only rods or both rods and cones (95% confidence interval, CI). C. Contact area with AIIs for RBCs contacting only rods or both rods and cones (95% CI). D. Dendritic (top) and axonal (bottom) mosaics for RBCs contacting rods or both rods and cones.

**Supplementary Figure 4:**
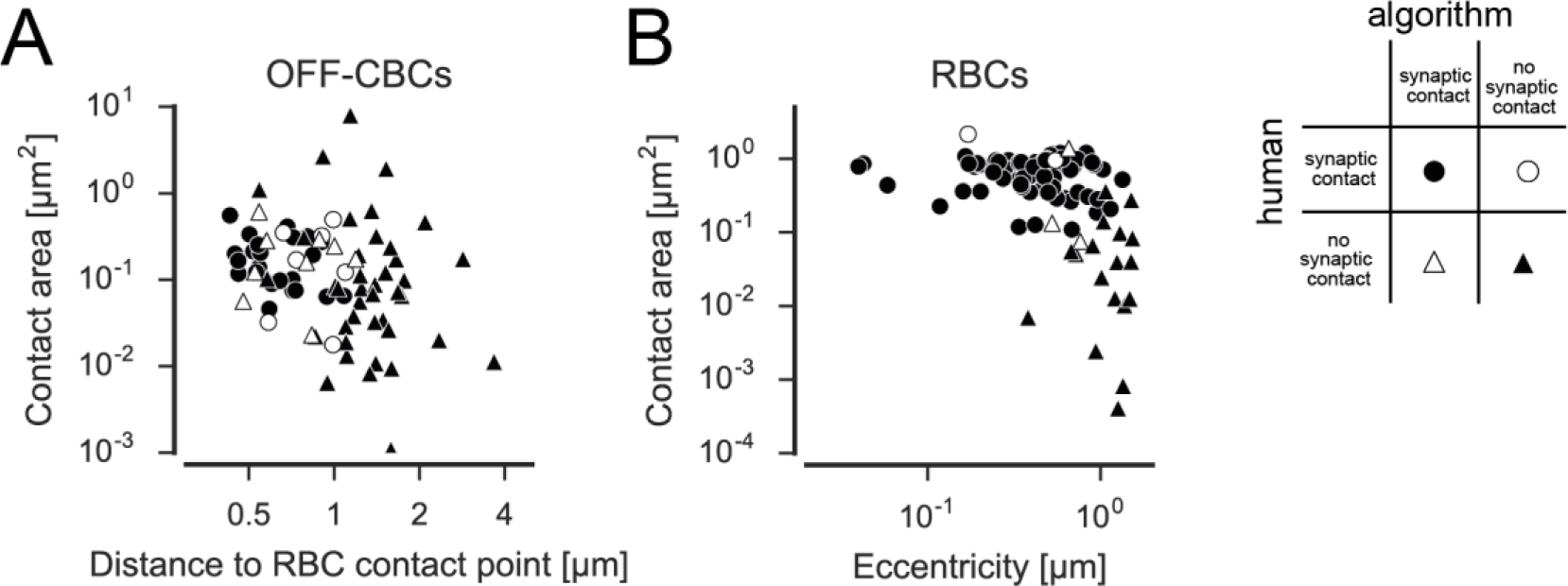
Contact area versus distance to RBC contact point for OFF-CBC-rod contacts (A) and contact area versus eccentricity for RBCs (B) contacts indicating correctly and incorrectly classified contacts.

**Supplementary Figure 5:**
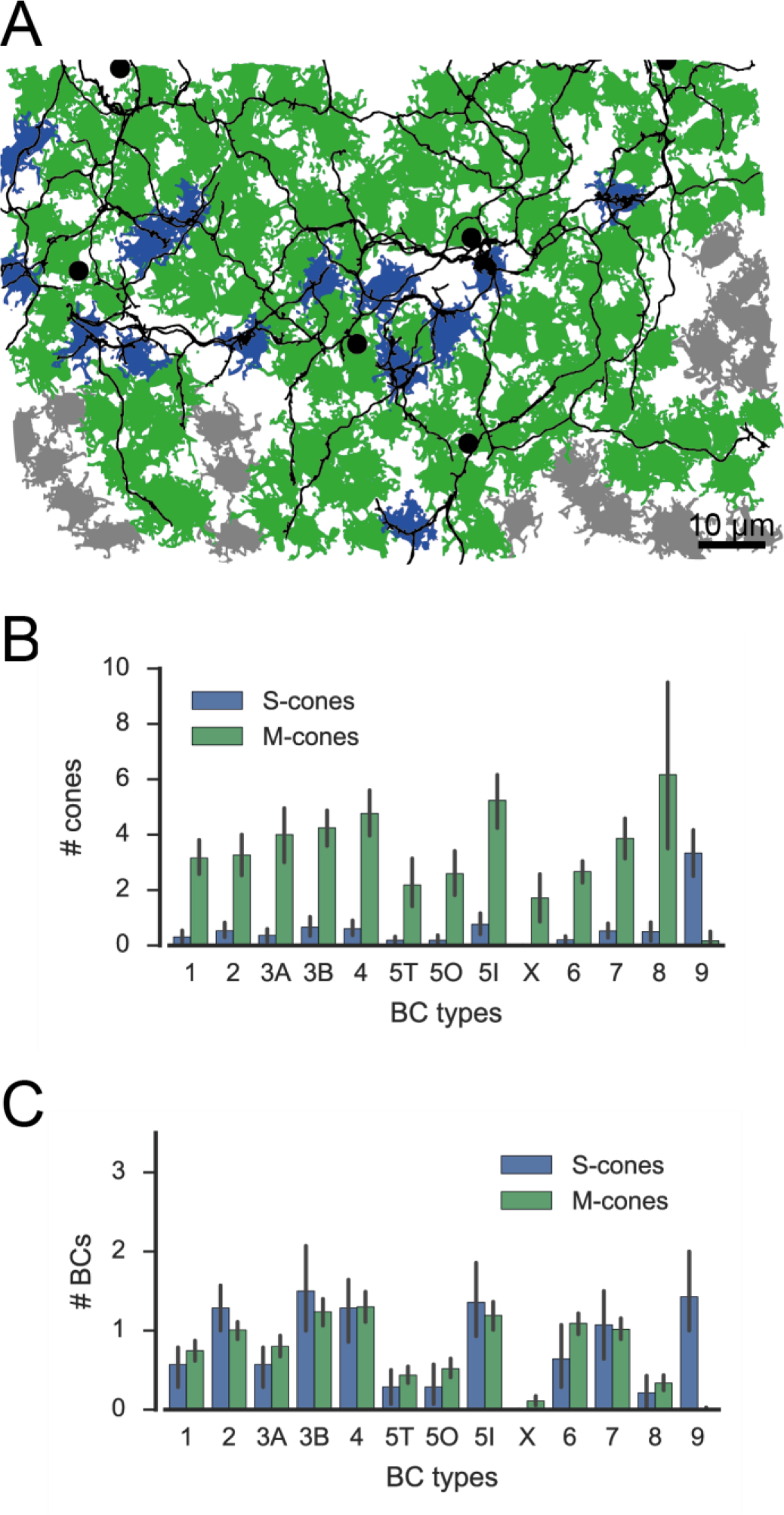
A. Cone pedicle array with CBC9s highlighted showing alternative S-cone classification. CBC9 somata are indicated by black dots, S-cones in blue, M-cones in green and unidentified cones in grey. B. Number of S-and M-cones contacted by different CBC types. C. Number of CBC types contacted by individual S-and M-cones.

**Supplementary Figure 6:**
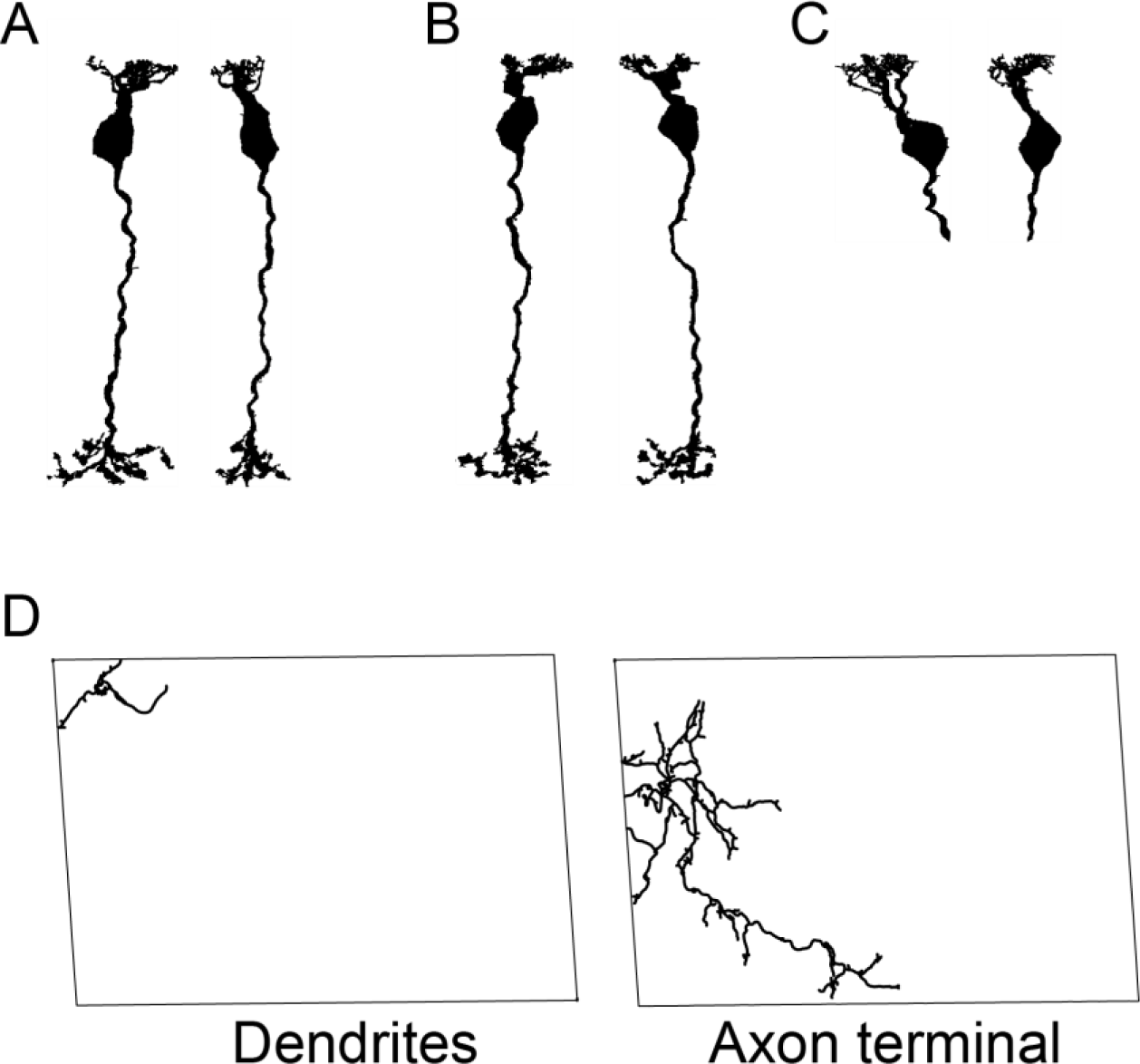
A-C. Three BCs classified as RBCs by Helmstaedter et al. (2013) but not contacting rods in the present study, these cells were therefore excluded from the analysis. D. BC classified as CBC9 but excluded from this study due to lack of complete dendritic tree.

**Supplementary Figure 7:**
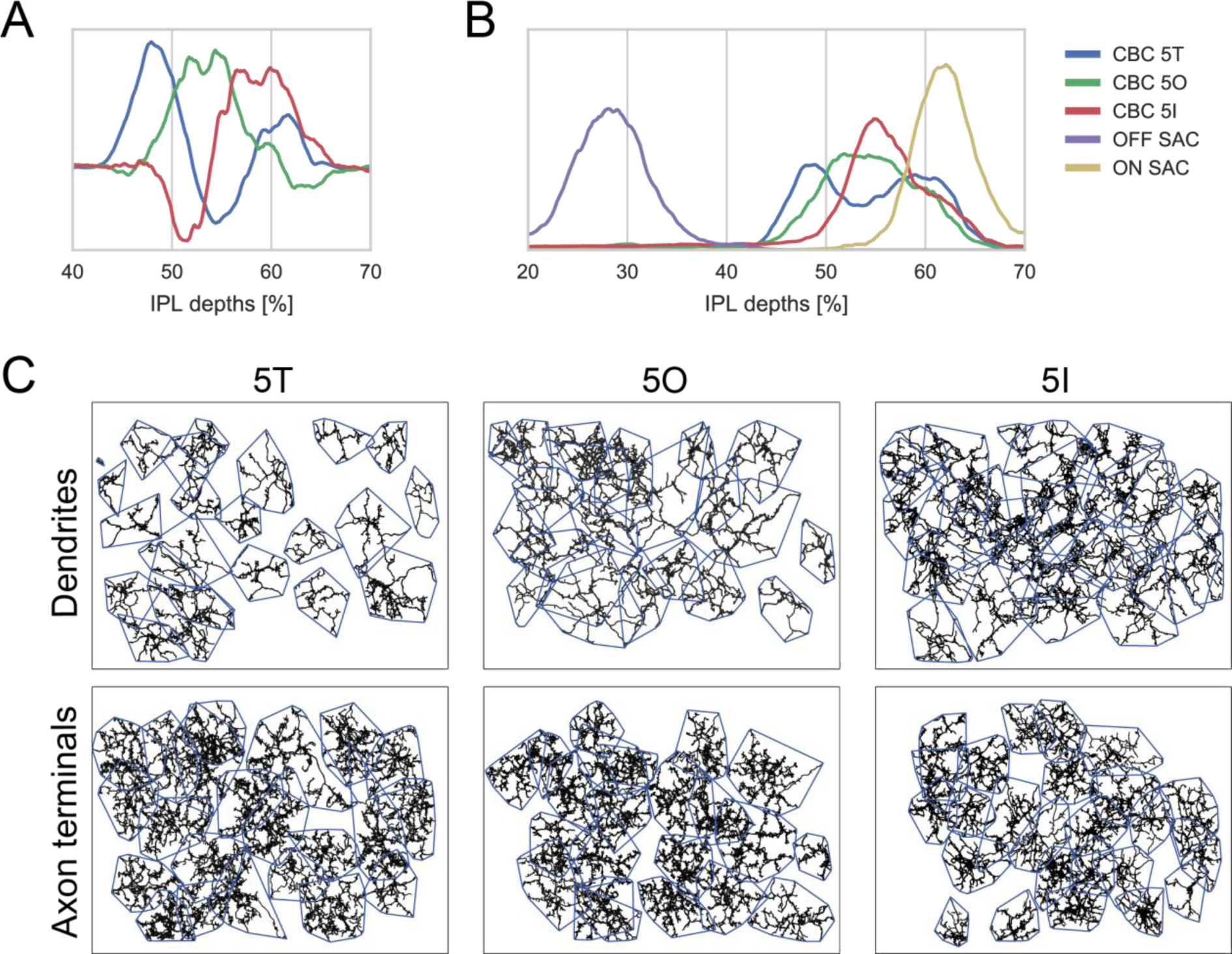
A. First three PCA components for CBC5 density profiles in the IPL. B. Stratification depth of CBC5T, 5O and 5I axon terminals in relation to the OFF-and ON-ChAT bands. C. Dendritic (top) and axonal (bottom) mosaics for CBC5T, 5O and 5I cells.

**Supplementary Figure 8:**
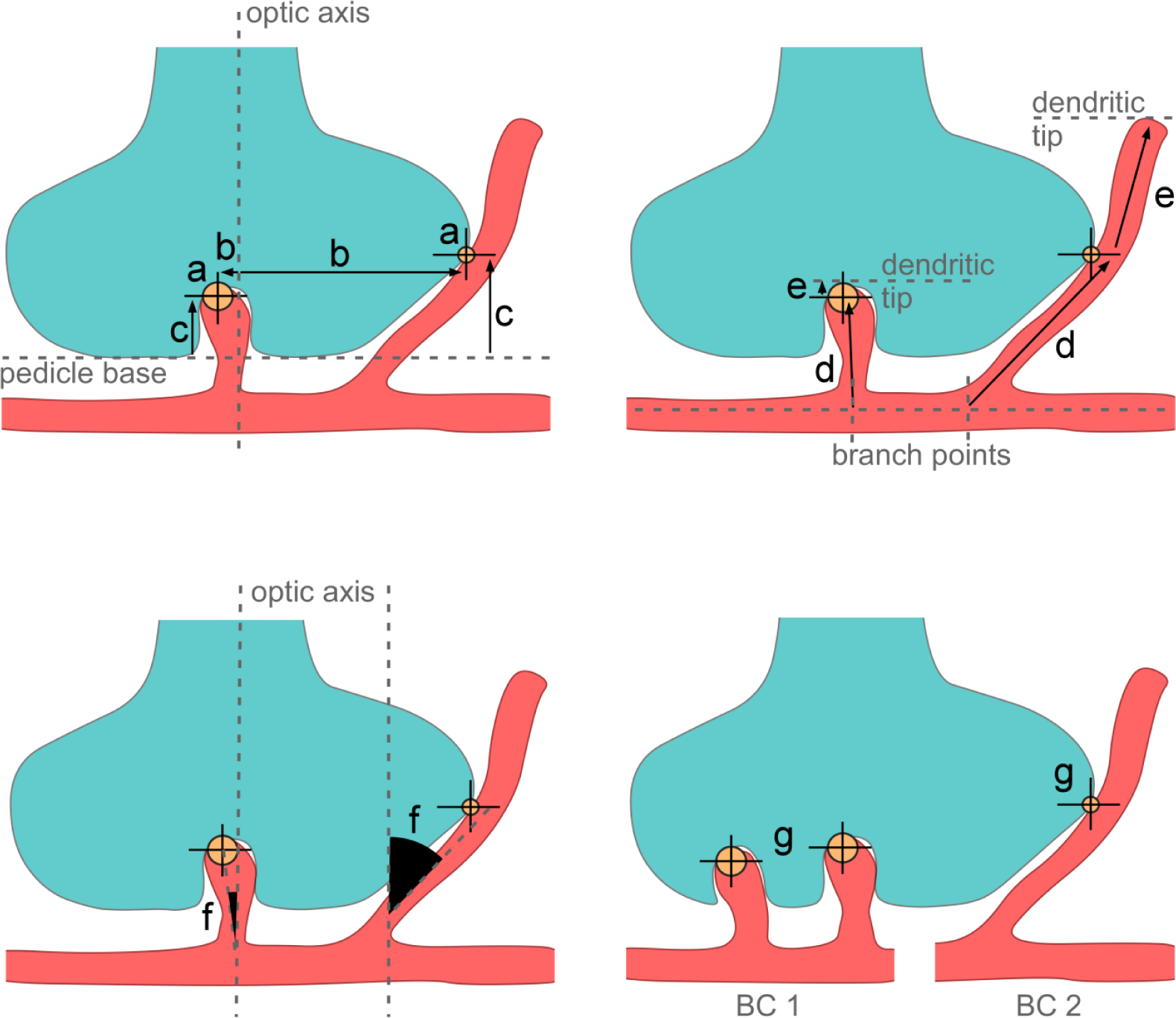
Cone pedicle schemes showing the parameter used for automated contact classification: Contact area (a), eccentricity (b), contact height (c), distances to branch point (d) and dendritic tip (e), smalles t angle between contacting dendrite and optical axis (g). Example invaginating and peripheral contacts between cone (cyan) and BC dedndrite(s) (red) are shown as large and small yellow circles, respectively. The optical axis is defined as a perpendicular through the centre of the cone pedicle. BC, bipolar cell.

## References

Baden T, Schubert T, Chang L, Wei T, Zaichuk M, Wissinger B, Euler T (2013) A tale of two retinal domains: Near-Optimal sampling of achromatic contrasts in natural scenes through asymmetric photoreceptor distribution. Neuron 80:1206–1217 Available at: http://dx.doi.org/10.1016/j.neuron.2013.09.030.

Bishop CM (2006) Pattern Recognition and Machine Learning. Springer New York.

Bloomfield S a, Xin D, Osborne T (1997) Light-induced modulation of coupling between All amacrine cells in the rabbit retina. Vis Neurosci 14:565–576.

Bloomfield SA, Dacheux RF (2001) Rod vision: Pathways and processing in the mammalian retina. Prog Retin Eye Res 20:351–384.

Boycott BB, Wässle H (1991) Morphological classification of bipolar cells of the primate retina. Eur J Neurosci 3:1069–1088.

Breuninger T, Puller C, Haverkamp S, Euler T (2011) Chromatic bipolar cell pathways in the mouse retina. J Neurosci 31:6504–6517.

Calkins DJ, Tsukamoto Y, Sterling P (1996) Foveal cones form basal as well as invaginating junctions with diffuse ON bipolar cells. Vision Res 36:3373–3381.

Chen M, Lee S, Park SJH, Looger LL, Zhou ZJ (2014) Receptive field properties of bipolar cell axon terminals in the direction-selective sublaminas of the mouse retina. J Neurophysiol Available at: http://www.ncbi.nlm.nih.gov/pubmed/25031256 [Accessed October 14, 2014].

Chun M-H, Grünert U, Martin PR, Wässle H (1996) The synaptic complex of cones in the fovea and in the periphery of the macaque monkey retina. Vision Res 36:3383–3395.

Della Santina L, Kuo SP, Yoshimatsu T, Okawa H, Suzuki SC, Hoon M, Tsuboyama K, Rieke F, Wong ROL (2016) Glutamatergic Monopolar Interneurons Provide a Novel Pathway of Excitation in the Mouse Retina.

DeVries SH, Li W, Saszik S (2006) Parallel Processing in Two Transmitter Microenvironments at the Cone Photoreceptor Synapse. Neuron 50:735–748.

Dowling JE., Boycott B. B. (1966) Organization of the Primate Retina: Electron Microscopy. Proc R Soc London, Ser B, Biol Sci 166:80–111.

Dunn F a., Wong ROL (2012) Diverse Strategies Engaged in Establishing Stereotypic Wiring Patterns among Neurons Sharing a Common Input at the Visual System’s First Synapse. J Neurosci 32:10306–10317.

Euler T, Haverkamp S, Schubert T, Baden T (2014) Retinal bipolar cells: elementary building blocks of vision. Nat Rev Neurosci 15:507–519 Available at: http://www.nature.com/doifinder/10.1038/nrn3783 [Accessed July 18, 2014].

Franke K, Berens P, Schubert T, Bethge M, Euler T, Baden T (2016) Balanced excitation and inhibition decorrelates visual feature representation in the mammalian inner retina. Available at: http://biorxiv.org/lookup/doi/10.1101/040642.

Greene MJ, Kim JS, Seung HS (2016) Analogous Convergence of Sustained and Transient Inputs in Parallel On and Off Pathways for Retinal Motion Computation. Cell Rep:1892–1900 Available at: http://linkinghub.elsevier.com/retrieve/pii/S2211124716300687.

Hack I, Peichl L, Brandstätter JH (1999) An alternative pathway for rod signals in the rodent 18 retina: rod photoreceptors, cone bipolar cells, and the localization of glutamate receptors. Proc Natl Acad Sci U S A 96:14130–14135.

Haverkamp S, Grünert U, Wässle H (2000) The Cone Pedicle, a Complex Synapse in the Retina. Neuron 27:85–95.

Haverkamp S, Specht D, Majumdar S, Zaidi NF, Brandstätter JH, Wasco W, Wässle H, Tom Dieck S (2008) Type 4 OFF cone bipolar cells of the mouse retina express calsenilin and contact cones as well as rods. J Comp Neurol 507:1087–1101.

Haverkamp S, Wässle H, Duebel J, Kuner T, Augustine GJ, Feng G, Euler T (2005) The primordial, blue-cone color system of the mouse retina. J Neurosci 25:5438–5445.

Helmstaedter M, Briggman KL, Denk W, Helmstaedter M, Briggman KL, Denk W (2012) High-accuracy neurite reconstruction for high-throughput neuroanatomy To cite this version:

Helmstaedter M, Briggman KL, Turaga SC, Jain V, Seung HS, Denk W (2013) Connectomic reconstruction of the inner plexiform layer in the mouse retina. Nature 500:168–174 Available at: http://www.nature.com/doifinder/10.1038/nature12346 [Accessed August 7, 2013].

Hopkins JM, Boycott BB (1995) Synapses between cones and diffuse bipolar cells of a primate retina. J Neurocytol 24:680–694.

Hopkins JM, Boycott BB (1996) The cone synapses of DB1 diffuse, DB6 diffuse and invaginating midget, bipolar cells of a primate retina. J Neurocytol 25:381–390.

Ichinose T, Fyk-Kolodziej B, Cohn J (2014) Roles of ON cone bipolar cell subtypes in temporal coding in the mouse retina. J Neurosci 34:8761–8771 Available at: http://www.jneurosci.org/content/34/26/8761.full [Accessed July 7, 2015].

Jiang X, Shen S, Cadwell CR, Berens P, Sinz F, Ecker AS, Patel S, Tolias AS (2015) Principles of connectivity among morphologically defined cell types in adult neocortex. Science (80-) 350:aac9462–aac9462 Available at: http://www.sciencemag.org/cgi/doi/10.1126/science.aac9462.

Joo HR, Peterson BB, Haun TJ, Dacey DM (2011) Characterization of a novel large-fieldcone bipolar cell type in the primate retina: evidence for selective cone connections. Vis Neurosci 28:29–37.

Kolb H (1970) Organization of the outer plexiform layer of the primate retina: electron microscopy of Golgi-impregnated cells. Philos Trans R Soc London B Biol Sci 258:261–283.

Kouyama N, Marshak DW (1992) Bipolar cells specific for blue cones in the macaque retina. J Neurosci 12:1233–1252.

Mariani AP (1984) Bipolar cells in monkey retina selective for the cones likely to be blue-sensitive. Nature 308:184–186 Available at: http://www.ncbi.nlm.nih.gov/htbin-post/Entrez/query?db=m&form=6&dopt=r&uid=6199677.

Mataruga A, Kremmer E, Müller F (2007) Type 3a and type 3b OFF cone bipolar cells provide for the alternative rod pathway in the mouse retina. J Comp Neurol 502:1123–1137 Available at: http://doi.wiley.com/10.1002/cne.21367.

Molnar A, Werblin F (2007) Inhibitory feedback shapes bipolar cell responses in the rabbit retina. J Neurophysiol 98:3423–3435 Available at:http://jn.physiology.org/content/jn/98/6/3423.full.pdf.

Pang J-J, Gao F, Lem J, Bramblett DE, Paul DL, Wu SM (2010)Direct rod input to cone BCs and direct cone input to rod BCs challenge the traditional view of mammalian BC circuitry. Proc Natl Acad Sci 107:395–400 Available at: http://www.pnas.org/cgi/doi/10.1073/pnas.0907178107.

Röhlich P, van Veen T, Szél Á (1994) Two different visual pigments in one retinal cone cell. Neuron 13:1159–1166 Available at: http://linkinghub.elsevier.com/retrieve/pii/0896627394900531.

Seung HS, Sümbül U (2014) Neuronal Cell Types and Connectivity: Lessons from the Retina. Neuron 83:1262–1272 Available at: http://linkinghub.elsevier.com/retrieve/pii/S0896627314007843 [Accessed September 17, 2014].

Szmajda B a., DeVries SH (2011) Glutamate Spillover between Mammalian Cone Photoreceptors. J Neurosci 31:13431–13441.

Tikidji-Hamburyan A, Reinhard K, Seitter H, Hovhannisyan A, Procyk CA, Allen AE, Schenk M, Lucas RJ, Munch TA (2015) Retinal output changes qualitatively with every change in ambient illuminance. Nat Neurosci 18:66–74 Available at: http://www.ncbi.nlm.nih.gov/pubmed/25485757.

Tsukamoto Y, Morigiwa K, Ishii M, Takao M, Iwatsuki K, Nakanishi S, Fukuda Y (2007) A novel connection between rods and ON cone bipolar cells revealed by ectopic metabotropic glutamate receptor 7 (mGluR7) in mGluR6-deficient mouse retinas. J Neurosci 27:6261–6267 Available at: http://www.jneurosci.org/cgi/doi/10.1523/JNEUROSCI.5646–06.2007 [Accessed July 20, 2016].

Tsukamoto Y, Omi N (2014) Some OFF bipolar cell types make contact with both rods and cones in macaque and mouse retinas. Front Neuroanat 8:105 Available at: http://www.ncbi.nlm.nih.gov/pubmed/25309346.

Wässle H, Puller C, Müller F, Haverkamp S (2009) Cone contacts, mosaics, and territories of bipolar cells in the mouse retina. J Neurosci 29:106–117.

